# CancerMine: A literature-mined resource for drivers, oncogenes and tumor suppressors in cancer

**DOI:** 10.1101/364406

**Authors:** Jake Lever, Eric Y. Zhao, Jasleen Grewal, Martin R. Jones, Steven J. M. Jones

## Abstract

Understanding a mutation in cancer requires knowledge of the different roles that genes play in cancer as drivers, oncogenes and tumor suppressors. We present CancerMine, a high-quality text-mined knowledgebase that catalogues over 856 genes as drivers, 2,421 as oncogenes and 2,037 as tumor suppressors in 426 cancer types. We compile 3,485 genes that are not in the IntOGen resource of drivers and complement the Cancer Gene Census with 3,136 new genes identified as oncogenes and tumor suppressors. CancerMine provides a method for gene-centric clustering of cancer types illustrating genetic similarities between cancer types of different organs and was validated against data from the Cancer Genome Atlas (TCGA) project. Finally with 178 novel cancer gene mentions in publications each month, this resource will be updated monthly, pre-empting the need to manually curate the ever-increasing number of novel cancer associated genes. CancerMine is viewable through a web portal (http://bionlp.bcgsc.ca/cancermine/) and available for download (https://github.com/jakelever/cancermine).

## Introduction

As sequencing technology becomes more widely integrated into clinical practice, genomic data from cancer samples is increasingly being used to support clinical decision making as part of precision medicine efforts. Many initatives use targeted panels that focus on well understood cancer genes, however more comprehensive approaches such as exome or whole genome sequencing that often uncover variants in genes of uncertain relevance to cancer are increasingly being employed. Interpreting individual cancer samples requires knowledge of which mutations are significant in cancer development. The importance of a particular mutation depends on the role of the associated gene and the specific cancer type. The terms “oncogene” and “tumor suppressor” are commonly used to denote genes (or aberrated forms) that respectively promote or inhibit the development of cancer. Genes of special significance to a particular cancer type or subtype are often described as “drivers”. A deletion or loss-of-function mutation in a tumor suppressor gene associated with the cancer type of the sample is potentially an important event for this cancer. Furthermore, amplifications and gain-of function mutations in oncogenes, and any somatic activity in known driver genes may be valuable information in understanding the mutational landscape of a given cancer sample. This knowledge can then help select therapeutic options and improve our understanding of markers of resistance in the particular cancer type.

A variety of methods exist to identify a gene as a driver, oncogene or tumor suppressor given a large set of genomic data. Many methods use the background mutation rate and gene lengths to calculate a p-value for the observed number of somatic events in a particular gene^1^. Other studies use the presence of recurrent somatic deletions or low expression to deduce that a gene is a tumor suppressor^2^. In-vitro studies that examine the effect of gene knockdowns on the cancer’s development are also used^3^.

Structured databases with information about the role of different genes in cancer, specifically as drivers, oncogenes and tumor suppressors, are necessary for automated analysis of patient cancer genomes. The Cancer Genome Atlas (TCGA) project has provided a wealth of information on the genomic landscape of over 30 types of primary cancers^4^. Data from TCGA (and other resources) are presented in the IntOGen resource to provide easy access to lists of driver genes^5^. The Cancer Gene Census has been curated using data from COSMIC to provide known oncogenes and tumor suppressors^6^ but faces the huge cost of manual curation. Other resources that provide curated information about cancer genes include TSGene^7^ and OnGene^8^ but do not match them with specific cancer types. There are also several other resources that are no longer accessible for unknown reasons (NCI Cancer Gene Index, MSKCC Cancer Genes database and Network of Cancer Genes^9^).

Text mining approaches can be used to automatically curate the role of genes in cancer, by identifying mentions of genes and cancer types, and extracting their relations from abstracts and full-text articles. Machine learning methods have shown great success in building protein protein interaction (PPI) networks using such data^10^. By weighting gene roles by the number of supporting papers and using a high-precision classifier, we mitigate the noisy biomedical corpora and extract highly relevant structured knowledge. We present CancerMine, a robust and regularly updated resource that describes drivers, oncogenes and tumor suppressors in all cancer types using the latest ontologies.

## Results

### Role of 3,775 unique genes catalogued in 426 cancer types

The entire PubMed, PubMed Central Open Access subset (PMCOA) and PubMed Central Author Manuscript Collection (PMCAMC) corpora were processed to identify sentences that discuss a gene and cancer types within titles, abstracts and where accessible full text articles. By filtering for a customized set of keywords, these sentences were enriched for those likely discussing the genes’ role and 1,500 randomly selected sentences were manually annotated by three expert annotators. Using a custom web interface and a well-defined annotation manual, the annotators tagged sentences that discussed one of three gene roles (driver, oncogene and tumor suppresser) with a mentioned type of cancer (Fig 1A). An example of a simple relation that was annotated as “Tumor Suppressor” annotation is: “DBC2 is a tumor suppressor gene linked to breast and lung cancers” (PMID: 17023000). A more complex example illustrates a negative relation: “KRAS mutations are frequent drivers of acquired resistance to cetuximab in colorectal cancers” (PMID:24065096). In this case, the KRAS mutations are drivers of drug resistance, and not of cancer development as required for annotation of driver relations.

**Figure 1.**
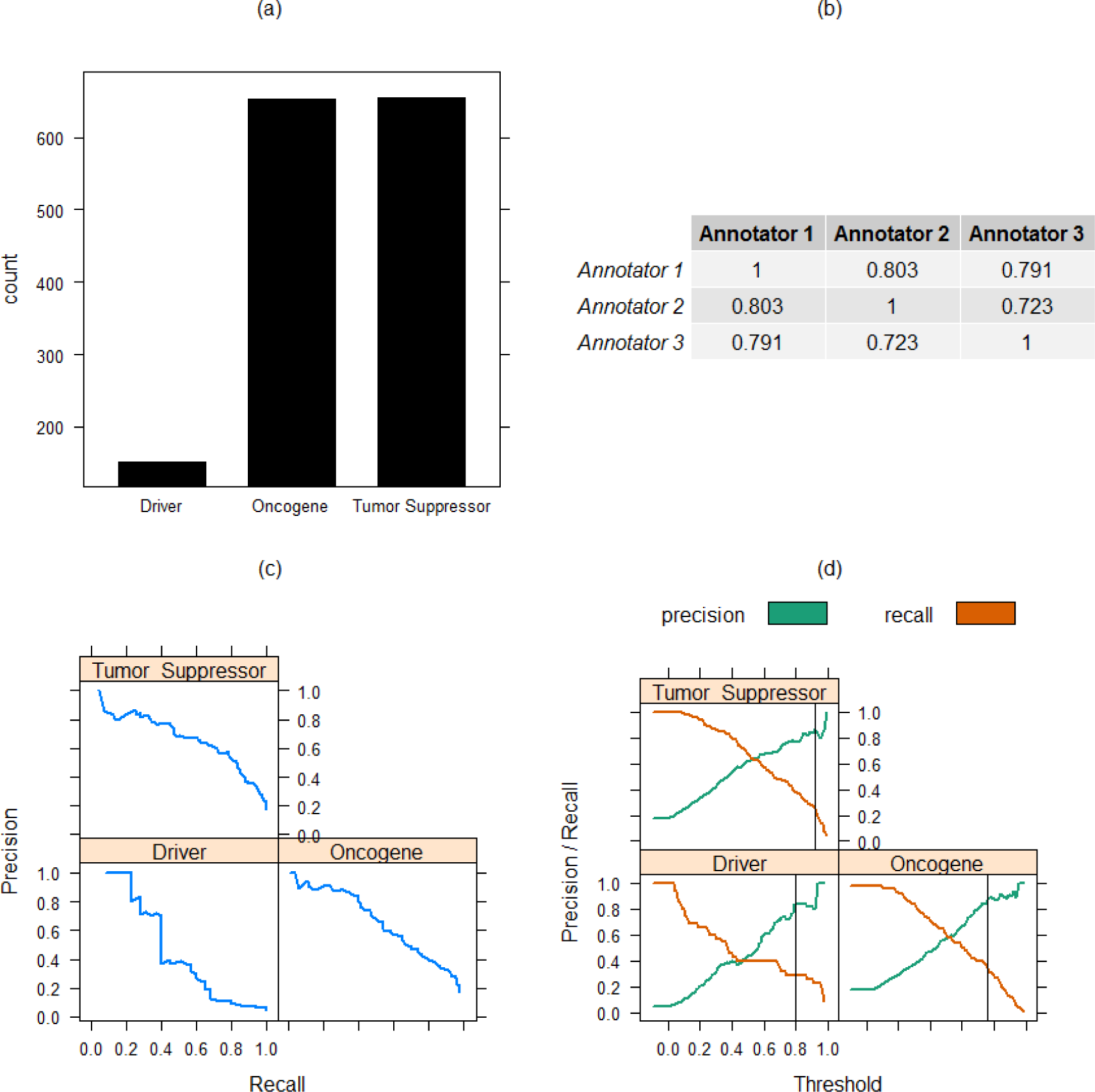
The supervised learning approach of CancerMine involves manual annotation by experts of sentences discussing cancer gene roles. Machine learning models are then trained and evaluated using this data set. (a) Manual text annotation of1,500 randomly selected sentences containing genes and cancer types show a similar number of Oncogene and Tumor Suppressor annotations. (b) The inter-annotator agreement (measured using Fl-score) was high between three expert annotators. (c) The precision recall curves show the trade-off of false positives versus false negatives. (d) Plotting the precision-recall data in relation to the threshold applied to the classifier’s decision function provides a way to select a high-precision threshold.

With high inter-annotator agreement (Fig 1B), the data were split into 75%/25% training and test sets. A machine learning model was built for each of the three roles and precision-recall curves were generated (Fig 1C) using the test set. Receiver operating characteristic (ROC) curves were not used as the class balance for each relation was below 20%. A high threshold was selected for each gene role in order to provide high-precision prediction with the accepted trade-off of low recall (Fig 1D).

The trade-off of higher precision with lower recall was made based on the hypothesis that there exists a large amount of redundancy within the published literature. The same idea is often stated multiple times in different papers in slightly different ways. Therefore, for frequently stated ideas, a method with lower recall would likely identify at least one occurrence. Nevertheless, we also distribute a version with thresholds of 0.5 for researchers who are willing to accept to a higher signal-to-noise ratio.

We apply the models to all sentences selected from PubMed abstracts and PMCOA/PMCAMC full-text articles, identifying 35,951 sentences from 26,767 unique papers that mention gene roles in cancer. We extract the unique gene/cancer pairs for each role (Fig 2A) and find that 3,775 genes and 426 cancer types are covered. These capture the commonly discussed cancer genes and types (Fig 2B/C) from a large variety of journals (Fig 2D). We provide a coverage of 21% (426/2,044) of the cancer types described in the Disease Ontology^11^ having at least one gene association. These results are made accessible through a web portal which can be explored through a gene or cancer-centric view. The resulting data are stored with Zenodo for versioning and download. This storage will provide the results in perpetuity. The results are licensed under the Creative Commons Public Domain (CC0) license to allow this data to be easily integrated with precision cancer workflows.

**Figure 2.**
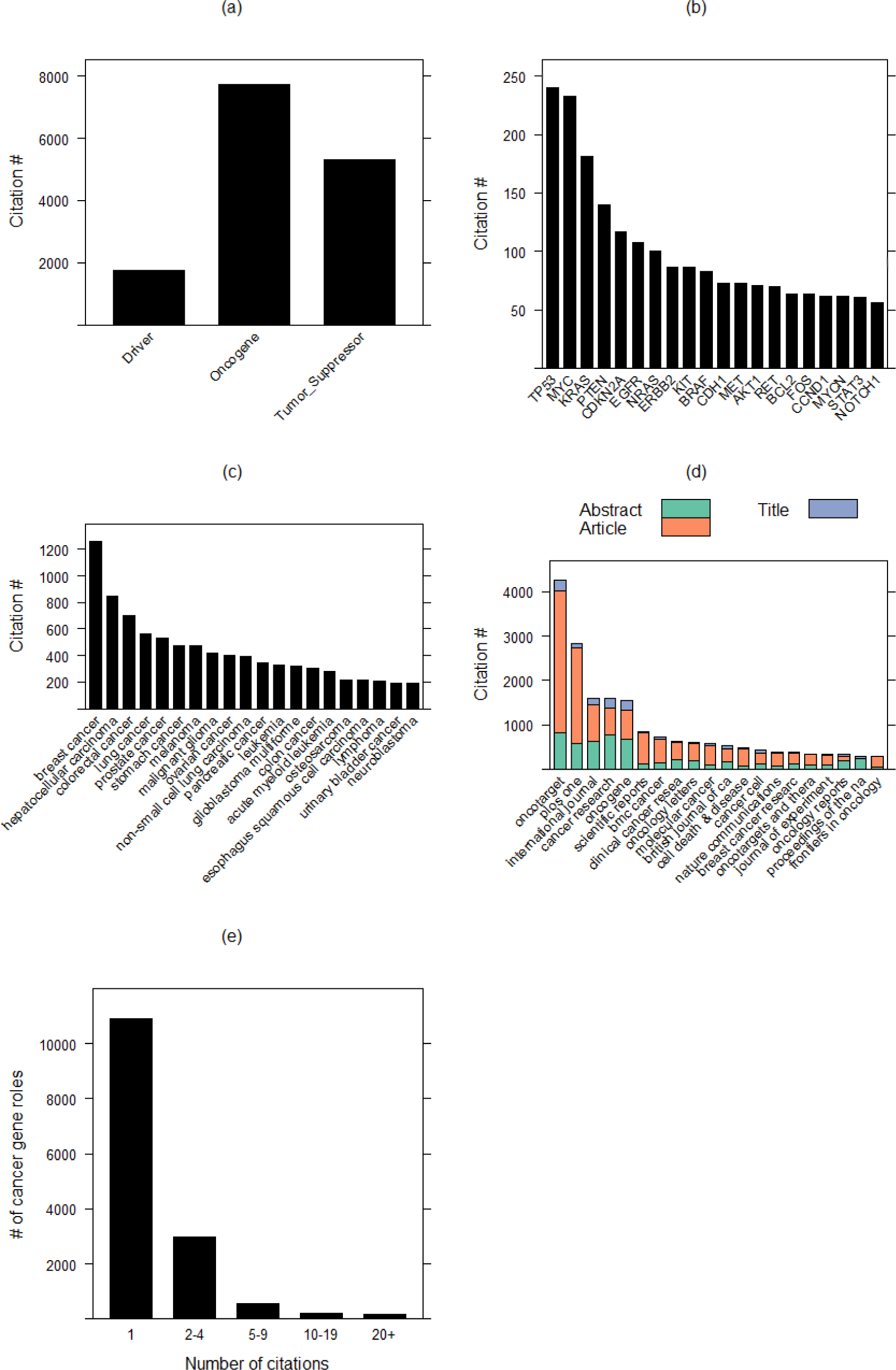
Overview of the cancer gene roles extracted from the complete corpora. (a) The counts of the three different gene roles extracted. (b) and (c) show the most frequently extracted genes and cancer types in cancer gene roles. (d) The most frequent journal sources for cancer gene roles with the section of the paper highlighted by color. (e) illustrates a large number of cancer gene roles have only a single citation supporting it but that a large number (3917) have multiple citations.

Our hypothesis of high levels of redundancy within the literature is supported by the frequent extraction of commonly-known gene roles such as ERBB2 as an oncogene in breast cancer (421 citations) and APC as a tumor suppressor in colorectal cancers (107 citations). On the other hand, a long tail exists of gene roles with only a single citation – 10,903 of 14,820 (73.6%) of extracted cancer gene roles (Fig 2E). For researchers that are accepting of a higher false positive rate, we provide an additional less stringent dataset using a lower prediction threshold and estimated average precision and recall of 0.5 and 0.6 respectively. The individual prediction score, akin to probabilities, are included so that users can further refine the results if needed.

### 60 novel putative tumor suppressors are published in literature each month

By examining the publication dates of the articles containing the mined cancer gene roles, we can see that the rate of published cancer gene roles is increasing over time (Fig 3A). In 2017, there were 6,851 mentions of cancer gene roles in publications, translating to over ~571 each month. Approximately 69% of these are gene roles that have been published previously, but more importantly, the remaining 31% are novel. A breakdown by the role shows that oncogene and tumor suppressor gene mentions greatly outnumber driver genes. In 2017, 1,358, 3,632 and 1,861 genes were mentioned as drivers, oncogenes and tumor suppressors (Fig 3B). Combining this data, we find that there were, on average, 22 novel drivers, 96 novel oncogenes and 60 novel tumor suppressors described in literature each month. This emphasizes the need to keep these text mining results up-to-date at a frequency of less than a year. To this end, we have integrated the CancerMine resource with the PubRunner framework to execute intelligent updates once a month (paper forthcoming - https://github.com/jakelever/pubrunner).

**Figure 3.**
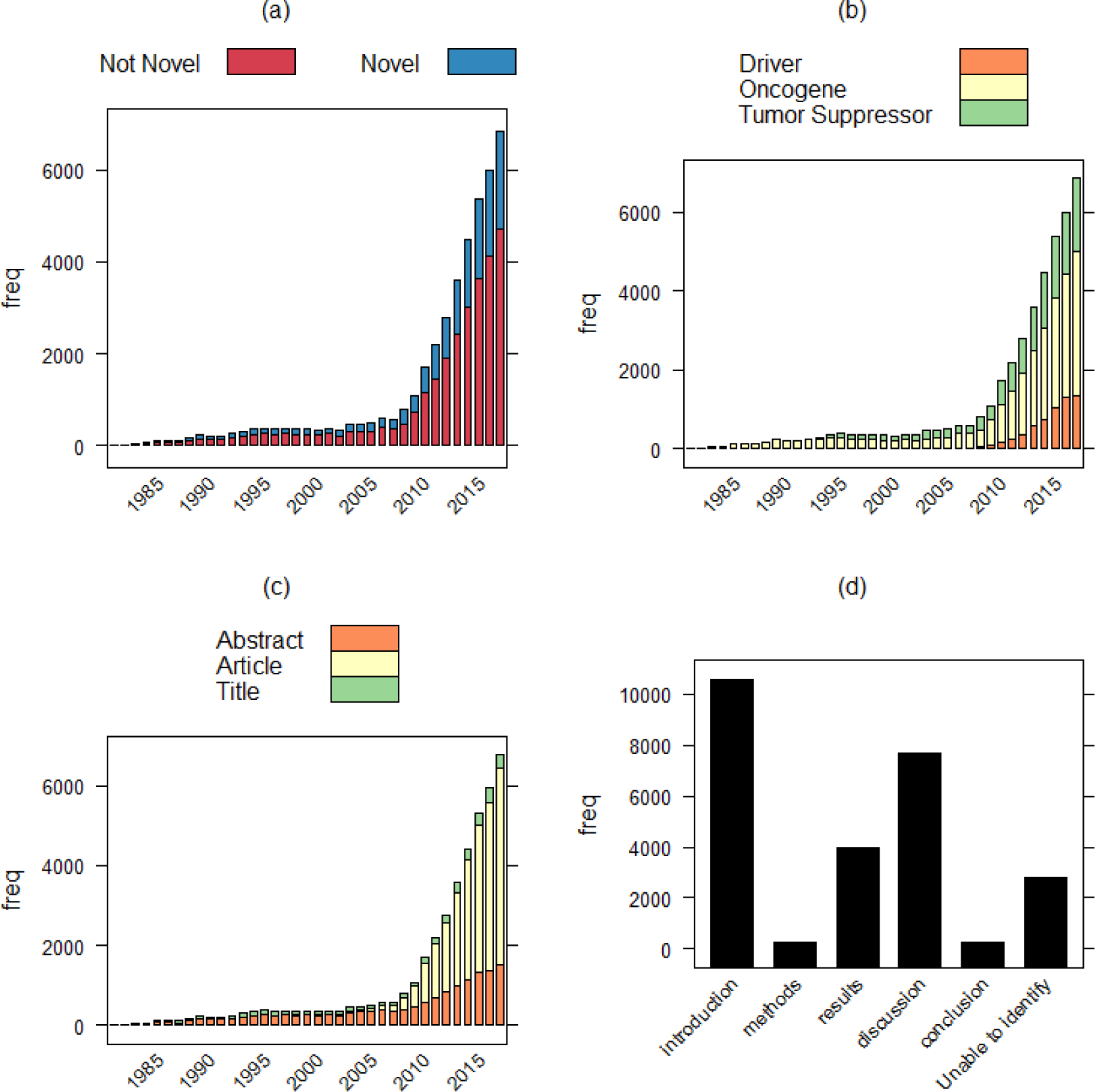
Examination of the sources of the extracted cancer gene roles with publication date. (a) More cancer gene roles are extracted each year but the relative proportion of novel roles remains roughly the same. (b) Roles extracted from older papers tend to focus on oncogenes, but mentions of driver genes have become more frequent since 2010. (c) The full text article is becoming a more important source of text mined data. (d) Different sections of the paper, particularly the Introduction and Discussion parts, are key sources of mentions of cancer gene roles (d).

Unhindered access to the full-text of articles for text mining purposes remains a key challenge. A larger number of cancer gene role mentions are extracted from the full text (25,641) than from the abstract alone (15,291), with a smaller number extracted from the titles (4,150). As can be seen in Fig 3C, the number extracted from full text articles is increasingly dramatically over time. This is likely linked to the increasing number of publications included in the PubMed Central Open Access subset and Author Manuscript Collection. This strengthens the need for publishers to provide open access and for funding agencies to require publications in platforms that allow text mining. From the full text articles, we extract, where possible, the in-text location of the relationship captured within the paper (Fig 3D). Interestingly, a substantial number of the mentions are found in the Introduction section, suggesting that the cancer gene’s role is usually discussed as background information and not a result of the paper. Knowing the subsection that a relationship is captured from can be valuable information for CancerMine users, since a user can then quickly ascertain if the discussed cancer role is prior knowledge or a likely result from the publication. This also highlights the important point that the scientific quality of a paper cannot be verified automatically by text mining technologies, since these methods rely on the statements made by the original author. Hence, any use of text-mined resources will always require users to access the original papers to evaluate the assertion of a gene’s role in a particular cancer.

Cancer gene roles that are first mentioned at earlier timepoints have more time to accrue additional citations (Fig 4A). Thus, it is no surprise that while most cancer gene roles have less than 10 associated citations, those with very large citation counts tend to be published over 10 years ago. For instance, ERBB2’s role as an oncogene in breast cancer is first extracted from a publication in 1988 and has accumulated 421 citations that fit our extraction criteria in literature since then. However, there are some cancer gene roles that were first extracted from publications within the last decade but have already accrued a great number of additional mentions. For instance, KRAS driving non-small cell lung carcinoma is first extracted from a paper published in 2010, and already has 92 other papers mentioning this role since. Lastly, there are 691 cancer gene roles that are mentioned in literature before 2000, but are not extracted in papers after that period. The most frequently mentioned cancer gene role that reflects this pattern is MYC as a oncogene in cervix carcinoma, with 10 papers mentioning it before 2000 but no further citations afterwards.

**Figure 4.**
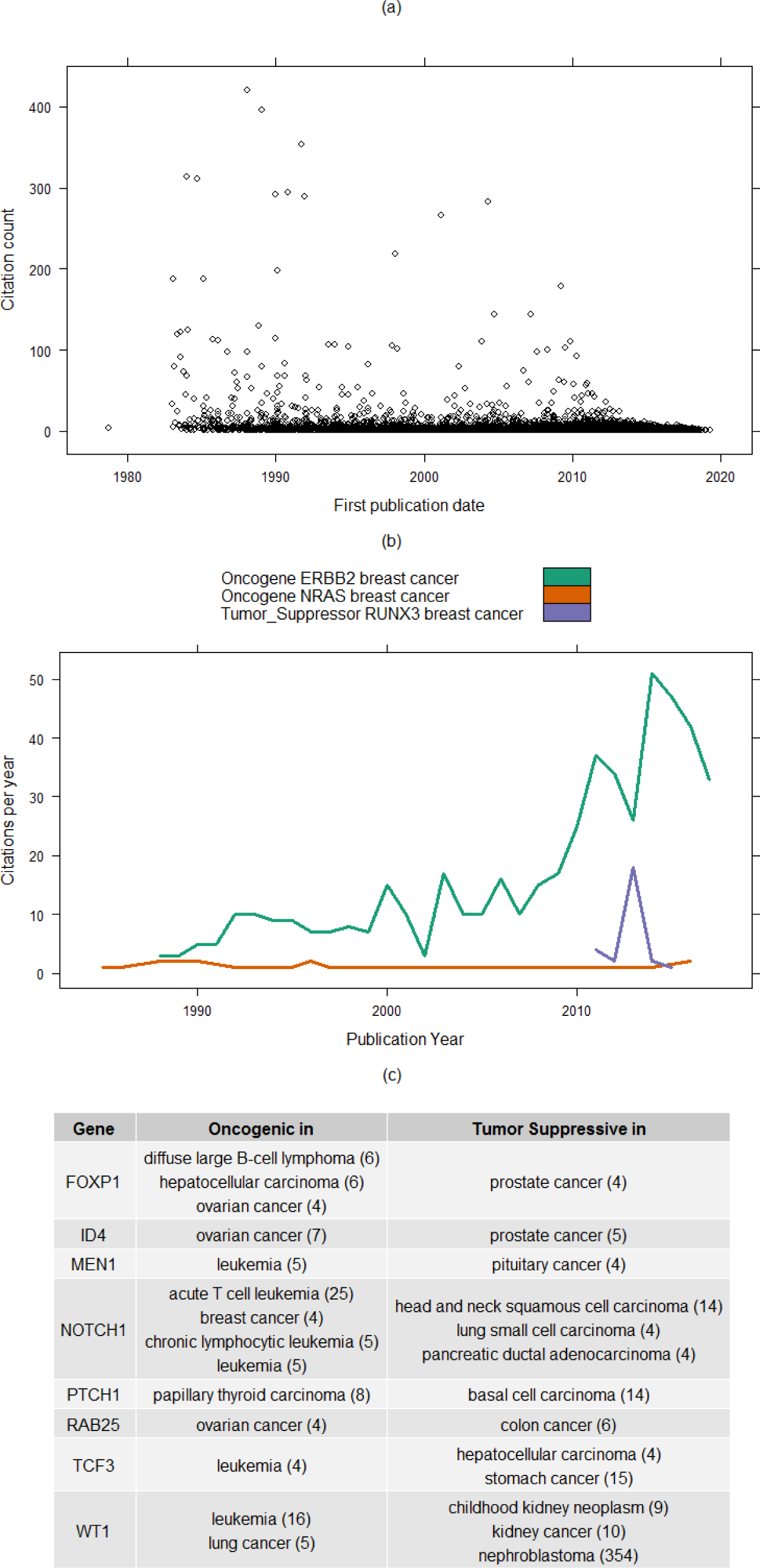
(a) Cancer gene roles discussed first discussed many years ago have a longer time to accrue further mentions. (b) Some cancer gene roles grow substantially in discussion while others fade away. (c) Cancermine can further validate the dual roles that some genes play as oncogenes and tumor suppressive. Citation counts are shown in parentheses.

With the knowledge of date of publication, we have gleaned a historical perspective on the gene relations captured in literature. Fig 4B summarizes three trends of citations that we observe, as exemplified by three gene associations with breast cancer. ERBB2 is an example of the small number of well established oncogenes that are more frequently discussed year upon year. NRAS in breast cancer exemplifies a gene that continues to be discussed in a single paper every few years, but has never gained importance in this cancer. RUNX3 has been discussed as a tumor suppressor in breast cancer in many papers in just the last few years. Its mechanism of action was elucidated after aggregated data from cell-line sequencing projects revealed its likely role as a tumor suppressor^12^.

The cancer type is important when trying to understand the context of somatic mutations. This is underscored by examples such as NOTCH1. NOTCH1 is a commonly-cited gene that behaves as an oncogene in one cancer (acute T cell leukemia) and as a tumor suppressor in another (head and neck squamous cell carcinoma)^13^. We further validate CancerMine by querying the resource for the set of genes that are (i) strongly identified as a oncogene in at least one cancer type (>90% of >=4 citations) and (ii) strongly identified as a tumor suppressor in at least one other cancer type. This method successfully identifies NOTCH1 along with several other genes that are reported to play dual roles in different cancer types (Fig 4C).

### Text mining provides voluminous complementary data to Cancer Gene Census

The Cancer Gene Census (CGC)^6^ provides manually curated information about cancer genes with mutation types and their roles in cancer. CancerMine contains information on 3,775 genes (compared to 554 in CGC) and 426 cancer types (compared to 201 in CGC). CancerMine overlaps with roughly a quarter of the oncogenes and tumor suppressors in the CGC when comparing specific cancer types (Fig 5A). When the CGC is compared to the less stringent CancerMine dataset, a further 202 cancer gene roles were found to match. This indicates that CGC contains curated information not easily captured using the sentence extraction method and that CancerMine represents an excellent complementary resource to work with CGC. Our resource also provides the sentence in which the gene role is discussed, and citations that link to the corresponding published literature are made available to help the user easily evaluate the evidence supporting the gene’s role. CancerMine would be an excellent resource for prioritizing future curation of literature for resources such as CGC.

**Figure 5.**
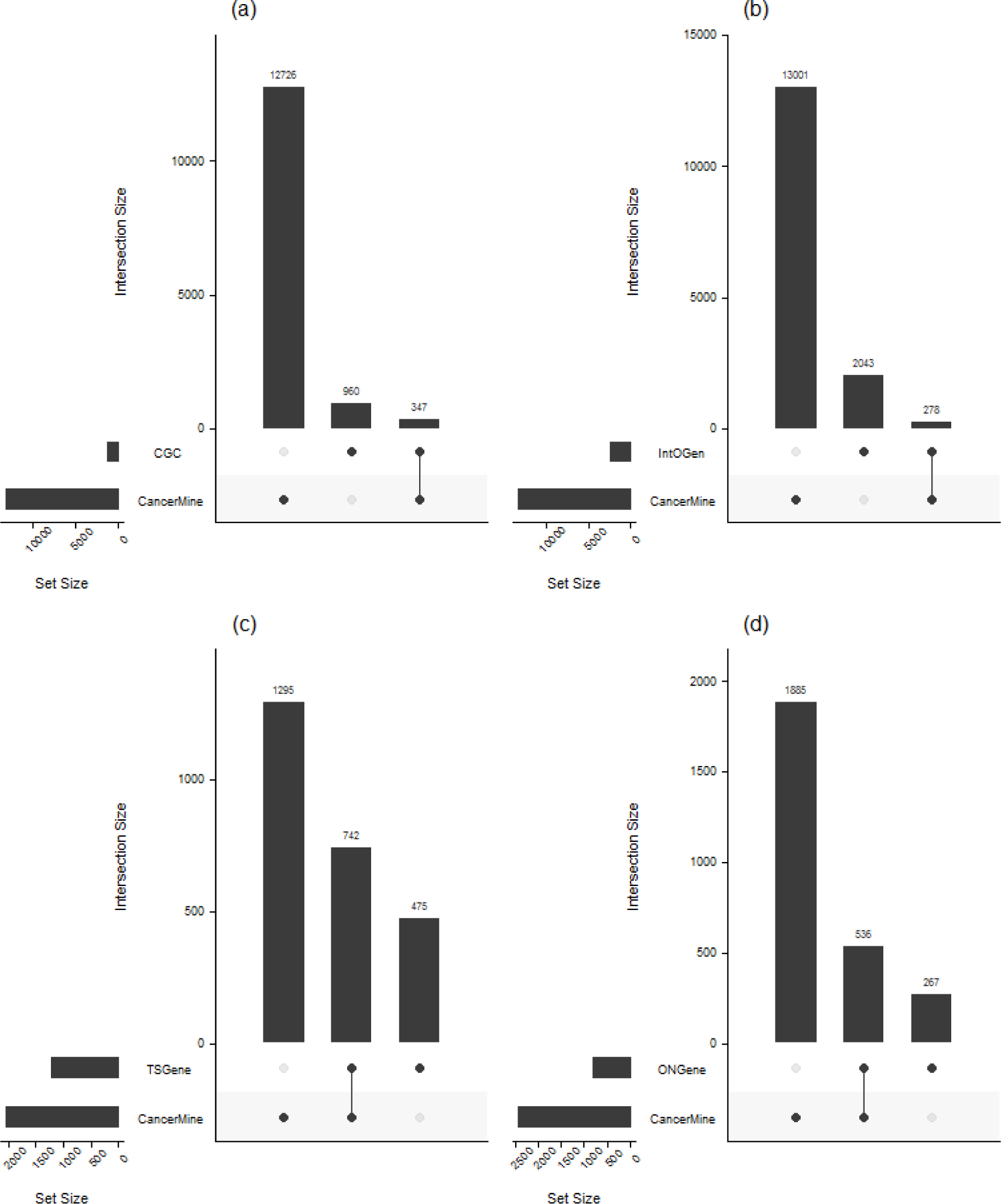
A comparison of CancerMine against resources that provide context for cancer genes. (a) The Cancermine resource contains substantially more cancer gene associations than the Cancer Gene Census resource. (b) Surprisingly few of the cancer gene associations are overlapping between the IntOGen resource and CancerMine. CancerMine overlaps substantially with the genes listed in the (c) TSGen and (d) ONGene resources.

The IntOGen resource leverages a number of cancer sequencing projects, including the Cancer Genome Atlas (TCGA) to index genes inferred to contain driver mutations. A comparison of the genes with their cancer types in Cancermine shows surprising differences (Fig 5B). CancerMine includes a much larger set of genes but has little overlap with the IntOGen resource. This suggests that many of the genes identified through the projects included in IntOGen are not yet frequently discussed in the literature with respect to the specific cancer types in the IntOGen resource.

ONGene and TSGene2 provide lists of oncogenes and tumor suppressors. Unfortunately these gene names are not associated with specific cancer types which is an important aspect for precision oncology. When trying to differentiate between driving and passenger mutations, the lack of cancer type context would likely cause a high false positive rate. CancerMine contains ~67% of the genes in ONGene and ~61% of TSGene2, and contains substantially more genes than both resources (Fig 5C/D). These results lend more weight to the use of automated text mining approaches for the population of knowledgebases, since no curation is required to keep the resource up-to-date.

Oncology often takes an organ-centric view of cancer types which is reflected by the numerous disease ontologies that exist for the categorization and nomenclature of cancer including the Disease Ontology used in this project. However, modern medicine is beginning to consider some cancers based purely on the genetic underpinnings, developing basket trials and approving treatment regimens based on genetic indications only (as shown with the successful approval of Pembrolizumab for PD-1 positive cancer patients). The CancerMine resource allows for the creation of a gene-centric view of cancers, by clustering cancers based on the role of different genes. A gene-centric view has the potential to reveal treatment regimes that could be transferred to other genetically similar cancer types. To allow for visualisation, we selected the top 30 cancers (based on citation count in CancerMine) and extracted the number of citations mentioning the role of the top 30 genes. This produces a profile for each cancer type showing the importance of each gene and its associated role. A heatmap that illustrates this for the top 30 cancer types and genes is shown in Figure 6a.

**Figure 6.**
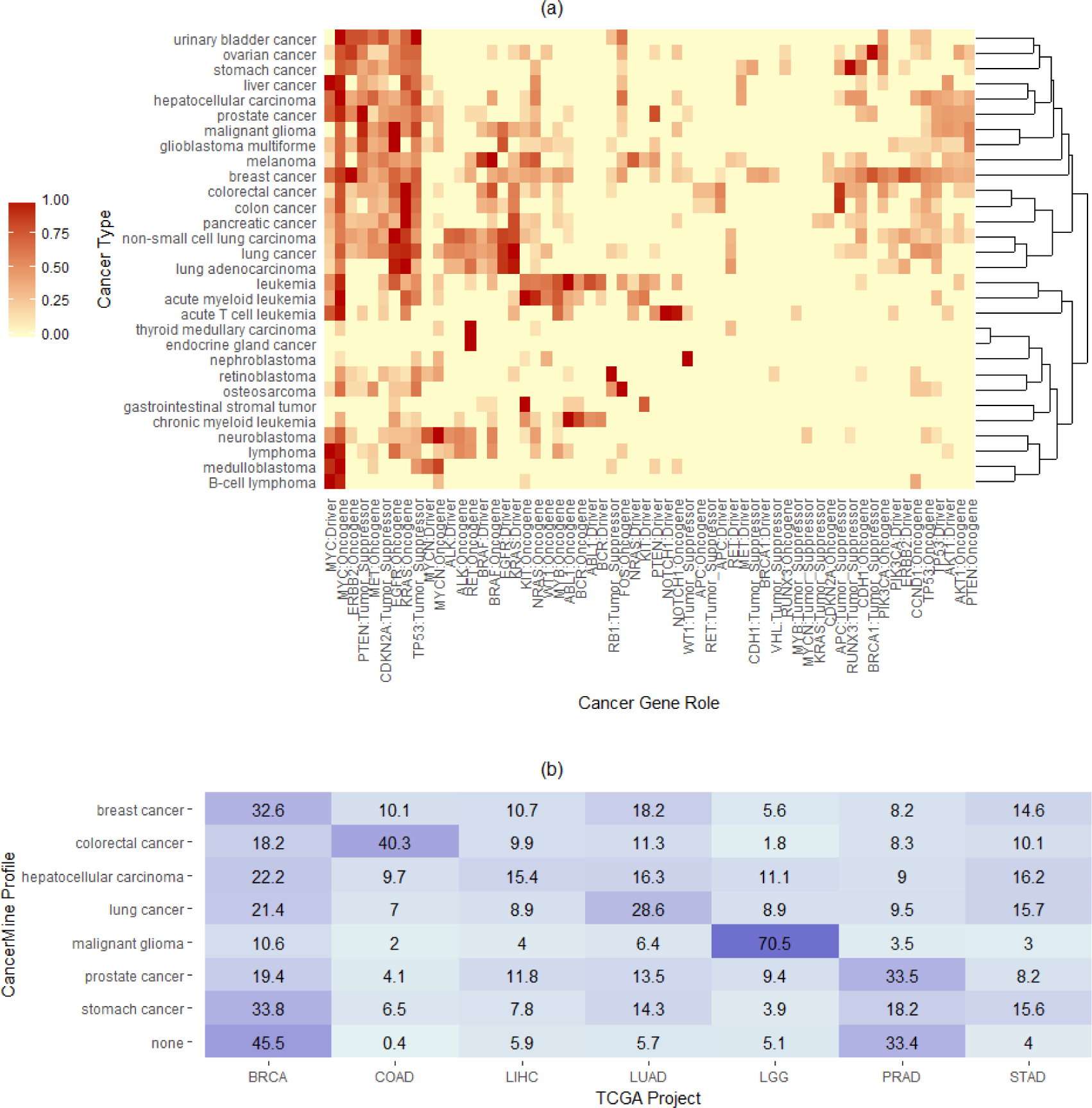
CancerMine data allows the creation of profiles for different cancer types using the number of citations as a weighting for each gene role. (a) The similarities between the top 30 cancer types in CancerMine are shown through hierarchical clustering of cancers types and genes using weights from the top 30 cancer gene roles. (b) All samples in seven TCGA projects are analysed for likely loss-of-function mutations compared with the CancerMine tumor suppressor profiles and matched with the closest profile. Percentages shown in each cell are the proportion of samples labelled with each CancerMine profile that are from the different TCGA projects. Samples that match no tumor suppressor in these profiles or are ambigious are assigned to none. The TCGA projects are breast cancer (BRCA), colorectal adenocarcinoma (COAD), liver hepatocellular carcinoma (LIHC), prostate adenocarcinoma (PRAD), low grade glioma (LGG), lung adenocarcinoma (LUAD) and stomach adenocarcinoma (STAD).

The clustering puts biologically similar or equivalent cancers together that are separate entities in the Disease Ontology. For example it groups colorectal with colon cancer and malignant glioma with glioblastoma multiforme. Some of these clusters also highlight known gene-cancer associations, for example, lung cancer, non-small cell lung carcinoma and lung adenocarinoma all cluster together, and are heavily associated with the KRAS and EGFR oncogenes. In fact, the strong cluster of genes on the left side separates cancers that are strongly associated with KRAS, EGFR, and TP53 (such as lung cancer) from those that are less so (such as thyroid medullary carcinoma). Put together, this approach is able to explain biological similarity of cancer types using shared gene associations. As an example, leukemia clusters closely with the more specific subtype, acute myeloid leukemia, and it is evident that this is driven by extracted associations of these cancers with MYC, ABL1 and many other genes. Several gene associations are noticeably low frequency compared to overall patterns, for instance KRAS in glioblastoma multiforme (GBM). While there are a small number of papers discussing KRAS in GBM, it is an infrequently discussed gene compared to EGFR and PTEN. Overall this visualisation presents an easy method to explore the similarities and differences between cancer types.

In order to validate the cancer genes identified in CancerMine, we compare results to somatic mutation data from the Cancer Genome Atlas (TCGA) project. We hypothesis that the genes denoted to be tumor suppressors would likely be affected by loss-of-function mutations. Using CancerMine profiles based on tumor suppressor genes, we compare somatic calls for all samples with mutation data within seven TCGA projects. For each sample, we match the somatic calls against the set of CancerMine tumor profiles and sum the importance of the tumor suppressors found to be mutated. Figure 6b shows the percentages of top matches to each CancerMine profile. Six of the seven CancerMine profiles have their highest proportion matches with the corresponding TCGA project. Interestingly a large number of breast cancer and prostate cancer samples cannot be unambigiously labelled with one of the CancerMine profiles. For prostate cancer, roughly one third of the samples do not have any LoF mutations that match against any tumor suppressors for any of the seven types, suggesting that prostate cancer tumor suppressors are disabled through other mechanisms or that there are more tumor suppressors involved which have not been captured by CancerMine.

The glioma (LGG) result is the most prominent with 70.5% of TCGA LGG samples being most closely identified with the CancerMine malignant glioma profile. This is largely due to the high prevalence of IDH1 (390/503) mutations identified in the LGG cohort. While this data would not be enough for a tumor type classifier on its own, this results shows there is substantial signal that can be leveraged for interpreting the genomic data and could be combined further with other mutational data. This is underscored when examining breast cancer tumor suppressors with only a single citation, genes that are hypothetically not well known to be tumor suppressors in breast cancer. Seven of these genes (ARID1B, FGFR2, KDM5B, SPEN, TBX3, PRKDC and KMT2C) are mutated in at least 10 TCGA BRCA samples providing extra strength for the importance of these genes in breast cancer. In fact, the mechanism through which KMT2C inactivation drives tumorgenesis was recently elaborated in ER-positive breast cancer^14^.

## Discussion

This work contributes a much needed resource of known drivers, oncogenes and tumor suppressors in a wide variety of cancer types. The text mining approach taken is able to discern complicated descriptions of cancer gene roles with a high level of precision. This provides for a continually updated resource with little need for human intervention. This generalizable method could extract other types of biological knowledge with only minor changes. However, there are several limitations to this approach that present interesting but challenging alleys for further investigation. Firstly, this method focuses on single sentence extraction due to the challenge of anaphora and coreference resolution across sentences. Previous work has shown the high false positive rate that frequently occurs when identifying knowledge across multiple sentences^15^. Our approach requires that authors discuss the gene name, role and cancer name all within the same sentence. This is a problem of writing style and probabilities that gets greatly diluted with the large number of publications processed. Furthermore our approach focuses on individual genes in isolation and is unable to capture complex interactions between cancer genes discussed in papers, e.g. mutual exclusivity. More of these complex relationships will likely be identified in future research and play a part in interpreting the somatic events in an individual cancer patient. Text mining approaches face growing challenges with extracting complex events like these, which may span multiple sentences or even paragraphs.

One important concept when interpreting CancerMine data is that our methodology does not force a definition of a driver, oncogene or tumor suppressor and relies on the assertion of individual authors. A decision was made to not extract discussion of genes frequently mutated in cancer. This was due to the acknowledged problem of huge genes (e.g. TTN) that frequently accrue many somatic mutations but likely don’t play a part in cancer. Instead we rely on the authors’ assertions of the role a gene plays in cancer. The level of evidence differs greatly as some assertions are based on intervential studies (e.g. knockdowns) while others use observational studies (e.g mutation frequency or expression experiments).

As has been noted, many attempts have been made to create a knowledgebase of this topic. Hosting the data through Zenodo and the code through Github provides a level of continuity that will guarantee that the project code and data stay accessible for the foreseeable future. Furthermore the PubRunner integration makes it easier to keep the results up-to-date. All data and analysis for this paper is open source and documented. We hope others will explore this data in order to infer new knowledge of cancer types and their associated genes.

## Online Methods

### Corpora Processing

PubMed abstracts and full-text articles from PubMed Central Open Access (PMCOA) subset and Author Manuscript Collection were downloaded using the PubRunner framework from the NCBI FTP websites. They were then converted to BioC format using PubRunner’s convert functionality. This strips out formatting tags and other metadata and retains the Unicode text of the title, abstract and for PMCOA, the full article. The source of the text (title, abstract, article) is also encoded.

### Entity recognition

Lists of cancer types and gene names were built using a subset of the Disease Ontology (DO) and NCBI gene lists. These were complemented by matching to the Unified Medical Language System (UMLS). For cancer types, this was achieved using the associated ID in DO or through exact string matching on the DO item title. For gene names, the Entrez ID was used to match with UMLS IDs. The cancer type was then associated with a DO ID, and the gene names were associated with their HUGO gene name. These cancer and gene lists were then pruned with a manual list of stop-words with several custom additions for alternate spellings/acronyms of cancers. All cancer terms with less than four letters were removed except for a selected set of abbreviations, e.g. GBM for glioblastoma multiforme.

The corpus text was loaded in BioC format and processed using the Kindred Python package which, as of v2.0, uses the Spacy IO parser. Using the tokenization, entities were identified through exact string matching against tokens. Longer entity names with more tokens were prioritised and removed from the sentence as entities were identified. Fusion terms (e.g. BCR-ABL1) were identified by finding gene names separated by a hyphen or slash. Non-fusions (e.g. HER2/neu) were then identified when two genes with matching HUGO IDs were attached and combined to be a single non-fusion gene entity. Genes mentioned in the context of pathways were also removed (e.g. MTOR pathway) using a list of pathway related keywords.

### Sentence selection

After Kindred parsing, the sentences with tagged entities were searched for those containing at least one cancer type and at least one gene name. These sentences were then filtered using the terms “tumor suppress”, “oncogen” and “driv” to enrich for sentences that were likely discussing these gene roles.

### Annotation

1,600 of the sentences were then randomly selected and output into the BioNLP Shared Task format for ingestion into an online annotation platform. This platform was then used by three expert annotators who are all PhD students actively engaged in precision cancer projects. The platform presents each possible pair of a gene and cancer and the user must annotate this as driving, oncogene and tumor suppressor. The first 100 sentences were used to help the users understand the system, evaluate initial inter-annotator agreement, and adjust the annotation guidelines (available at the Github repository). The results were then discarded and the complete 1,500 sentences were annotated by the first two annotators. The third annotator then annotated the sentences that the first two disagreed on. The inter-annotator agreement was calculated using the F1-score. A gold corpus was created using the majority vote of the annotations of the three annotators.

### Relation extraction

75% of the 1500 sentences were used as a training set and a Kindred relation classifier was trained with an underlying logistic regression model for all three gene roles (Driver, Oncogene and Tumor_Suppressor). The threshold was varied to generate the precision-recall curves with evaluation on the remaining 25% of sentences. With the selection of the optimal thresholds, a complete model was trained using all 1,500 sentences. This model was then applied to all sentences found in PubMed, PMCOA and PMCAMC that fit the sentence requirements. The associated gene and cancer type IDs were extracted, entity names were normalized and the specific sentence was extracted.

### Web portal

The resulting cancer gene roles data were aggregated by the triples (gene, cancer, role) in order to count the number of citations supporting each cancer gene role. This information was then presented through tabular and chart form using a Shiny web application.

### Resource comparisons

The data from the Cancer Gene Census (CGC), IntOGen, TS and ONGene resources were downloaded for comparison. HUGO gene IDs in CancerMine were mapped to Entrez gene IDs. CGC data was mapped to Disease Ontology cancer types using a combination of the cancer synonym list created for CancerMine and manual curation. Oncogenes and tumor suppressors were extracted using the presence of “oncogene” or “TSG” in the “Role in Cancer” column. The mapped CGC data was then compared against the set of oncogenes and tumor suppressors in CancerMine. IntOGen cancer types were manually mapped to corresponding Disease Ontology cancer types and compared against all of CancerMine. The TSGene and ONGene gene sets were compared against the CancerMine gene sets without an associated cancer type.

### CancerMine profiles and TCGA analysis

For each cancer type, the citation counts for each gene role that were in the top 30 cancer genes were then log10-transformed and rescaled so that the most important gene had the value of 1 for each cancer type. Gene roles with values lower than 0.2 for all cancer types were trimmed. The top 30 cancer types and genes were then hierarchical clustered for the associated heatmap.

The open-access VarScan somatic mutation calls for the seven TCGA projects (BRCA,COAD,LIHC,PRAD,LGG,LUAD,STAD) were downloaded from the GDC Data Portal (https://portal.gdc.cancer.gov). They were filtered for mutations that contained a stop gain or were classified as probably damaging or deleterious by PolyPhen. Tumor suppressor specific CancerMine profiles were generated that used all tumor suppressors for each cancer type. The citation counts were again log10-transformed and rescaled to produce the CancerMine tumor suppressor profile. Each TCGA sample was represented as a binary vector matching the filtered mutations. The dot-product of a sample vector and a CancerMine profile vector produced the sum of citation weightings and gave the score. For each sample, the score was calculated for all seven cancer types and the highest score was used to label the sample. A sample that did not contain tumor suppressor mutations associated with any of the seven profiles or could not be labelled unambigously was labelled as ‘none’.

## Acknowledgements

JL was supported by a Vanier Canada Graduate Scholarship. The authors would like to thank Compute Canada for the computational resources used in this work.

## Author Contributions

JL, MRJ and SJMJ developed the idea for the project. MRJ and SJMJ supervised the project. JL, EYZ and JG annotated training data for the project. JL developed and implemented all methods for this project. All authors contributed to the manuscript.

